# Probiotic Lactobacilli activate Formyl-Peptide Receptor 2

**DOI:** 10.1101/2024.05.07.592932

**Authors:** Kretschmer Dorothee, Rosenstein Ralf, Elsherbini Ahmed, Krismer Bernhard, Paul W. O’Toole, Gerlach David, Peschel Andreas

## Abstract

Changes in the composition of the human microbiota can negatively impact human health. Probiotic bacteria like many lactobacilli help prevent or repair dysbiosis but it is largely unclear which molecules of these bacteria mediate the probiotic effects. Given the extensive crosstalk between the immune system and microbiome members, we investigated whether lactobacilli activate the formyl-peptide receptor 2 (FPR2), a pattern recognition receptor that is expressed on the surface of intestinal epithelial cells and known to promote wound healing and immune homeostasis.

Probiotic strains of *Lacticaseibacillus paracasei, Lactiplantibacillus plantarum*, and *Lacticaseibacillus rhamnosus* were isolated from probiotic compounds and sequenced. Calcium influx experiments in FPR1 or FPR2 overexpressing HL60 cells, and primary human neutrophils, along with pharmacological inhibition of FPR2, revealed that culture filtrates of the isolated lactobacilli strongly activate FPR2, promote killing of the methicillin resistant *S. aureus* USA300 and induce neutrophil chemotaxis. Pretreatment of culture filtrates with proteinase K reduced FPR2 activity, indicating that the FPR2 ligands are peptides. In silico analysis of the amphipathic properties of the signal peptides of lactic acid bacteria identified selected signal peptides of *L. plantarum* with the ability to predominantly activate FPR2 *in vitro*. Thereby, via targeted activation of FPR2, peptides released by some lactobacilli are likely to positively influence the outcome of inflammatory gut diseases and could be used to treat inflammatory diseases.

## Introduction

Pathogenic and commensal bacteria have many properties in common. They release microbe-associated molecular patterns (MAMPs), which are either metabolites or cell components, e.g., flagellin, with the ability to activate different pattern recognition receptors (PRRs) of the host (Clasen et al. 2023). N-terminal formylation is a hallmark of bacterial peptides as only prokaryotic ribosomes start protein biosynthesis with a formylated methionine (Schiffmann, Corcoran, and Wahl 1975). Nevertheless, bacteria differ in the amounts of formyl groups that are cleaved off by deformylases (Yuan and White 2006). They also differ in the type and amounts of signal peptides that are cleaved off from membrane or secreted proteins by dedicated peptidases (de Souza et al. 2011; Ravipaty and Reilly 2010; Bufe et al. 2015). Accordingly, host cells are equipped to respond to different types of bacterial peptides via a unique class of PRRs, the so-called formyl-peptide receptors (FPRs). While FPR1 exclusively senses short, formylated, hydrophobic peptides, FPR2 accepts longer, α-helical, amphipathic peptides (Kretschmer et al. 2015; Forsman et al. 2015). FPR2 prefers formylated peptides too (Rautenberg et al. 2011), but responds also to some non-formylated peptides including certain human peptides such as the antimicrobial peptide LL-37 (Singh et al. 2013). Virtually all bacteria activate FPR1, but it is unclear how widespread the ability to activate FPR2 is among bacteria. We have found that many staphylococci and some enterococci activate FPR2, but whether FPR2 activation is a widespread trait and plays a role in probiotic bacteria is still unclear.

The human microbiota, especially the one in the gut, is crucial for human health and changes in its composition, also known as dysbiosis, are associated with different pathological states (Beller et al. 2021; Talapko et al. 2022). Probiotic bacteria can help avoid or repair dysbiosis and a few probiotic bacteria have been described for the treatment of diarrhea (Hou et al. 2020), but the reason for their probiotic effects are often undefined. While intestinal bacteria are known to communicate with gut epithelia cells (Kaur, Ali, and Yan 2022), it remains unclear which bacterial agonists distinguish probiotic from non-probiotic bacteria.

Gut epithelia cells possess a variety of receptors to recognize bacterial molecules such as peptidoglycan, lipopeptides, or formylated peptides (Burgueno and Abreu 2020; Chen et al. 2023; Alam et al. 2014). Among those receptors, FPR2 has a special role. It is unresponsive to most bacterial species except for intestinal enterococci, some of which are used as therapeutic probiotics and produce FPR2 agonists (Bloes et al. 2012). In addition, skin-colonizing staphylococci can activate leucocytes via FPR2. FPR2 activation of intestinal epithelia cells induces ROS production and thereby leads to improved wound healing and immune homeostasis (Birkl et al. 2019). Interestingly, FPR2 expression is upregulated in inflamed intestinal tissue of patients with Crohn’s disease (Prescott and McKay 2011). Moreover, endogenous FPR2 ligands such as lipoxin A4 or annexin positively influence the course of disease by dampening inflammation via FPR2 (Vong et al. 2012).

The best known and most potent stimulators of FPR2 among the bacteria belong to the Firmicutes. Potentially probiotic bacteria like lactobacilli also belong to the Firmicutes and promote immune homeostasis in the gut. We asked whether they can also activate FPR2. We found that culture filtrates of lactic acid bacteria, including *Lacticaseibacillus paracasei, Lactiplantibacillus plantarum* and *Lacticaseibacillus rhamnosus*, strongly activate FPR2 and to a minor degree FPR1 on the surface of FPR1- or FPR2-overexpressing HL60 cells and primary human neutrophils. Digestion with proteinase K and cultivation with the deformylase inhibitor actinonin revealed that the FPR2 ligands are formylated peptides. It was shown that signal peptides from many bacteria could represent a pool of formylated peptides. Therefore, we speculated that the FPR2 ligands in culture filtrates of lactobacilli could be signal peptides. To investigate this, we sequenced the genome of *L. paracasei, L. plantarum* and *L. rhamnosus* and explored *in silico* the amphipathic properties of the signal peptides of these and further lactic acid bacteria. The amphipathic properties of some signal peptides fitted into this assumption as observed for α-PSMs, known FPR2 ligands. To experimentally prove this, we analyzed selected signal peptides of *L. plantarum* regarding their FPR2 activity and observed that the analyzed signal peptides of *L. plantarum* strongly activate FPR2. Our results indicate that lactobacilli release signal peptides that activate predominantly FPR2. Thereby, they are likely to positively influence the outcome of inflammatory gut diseases.

## Results

### Culture filtrates of lactobacilli activate formyl-peptide receptors

We analyzed culture filtrates of various bacteria like *Escherichia coli, Yersinia pseudotuberculosis, Streptococcus agalactiae, Enterococcus faecium* and *E. faecalis* for their ability to activate FPR1 and FPR2. Only for the culture filtrates of *Enterococcus faecium* and *Lacticaseibacillus rhamnosus* we observed a moderate and a strong FPR2 activity, respectively, measured as calcium influx induction (Fig.1 A). To assess whether lactic acid bacteria can activate FPRs, we tested whether culture filtrates of various lactobacilli induced calcium influx in FPR1- or FPR2-overexpressing HL60 cells. We observed that culture filtrates of these lactobacilli strains activated to a moderate degree FPR1, and that diluted culture filtrates of some of them *(L. paracasei, L. plantarum, L. rhamnosus*) activated FPR2. Interestingly, all analyzed culture filtrates also induced calcium influx in primary human neutrophils, which express FPR1 and FPR2. Accordingly, preincubation of neutrophils with the FPR2 inhibitor WRW4 significantly reduced activation of PMNs by culture filtrates.

**Figure 1.**
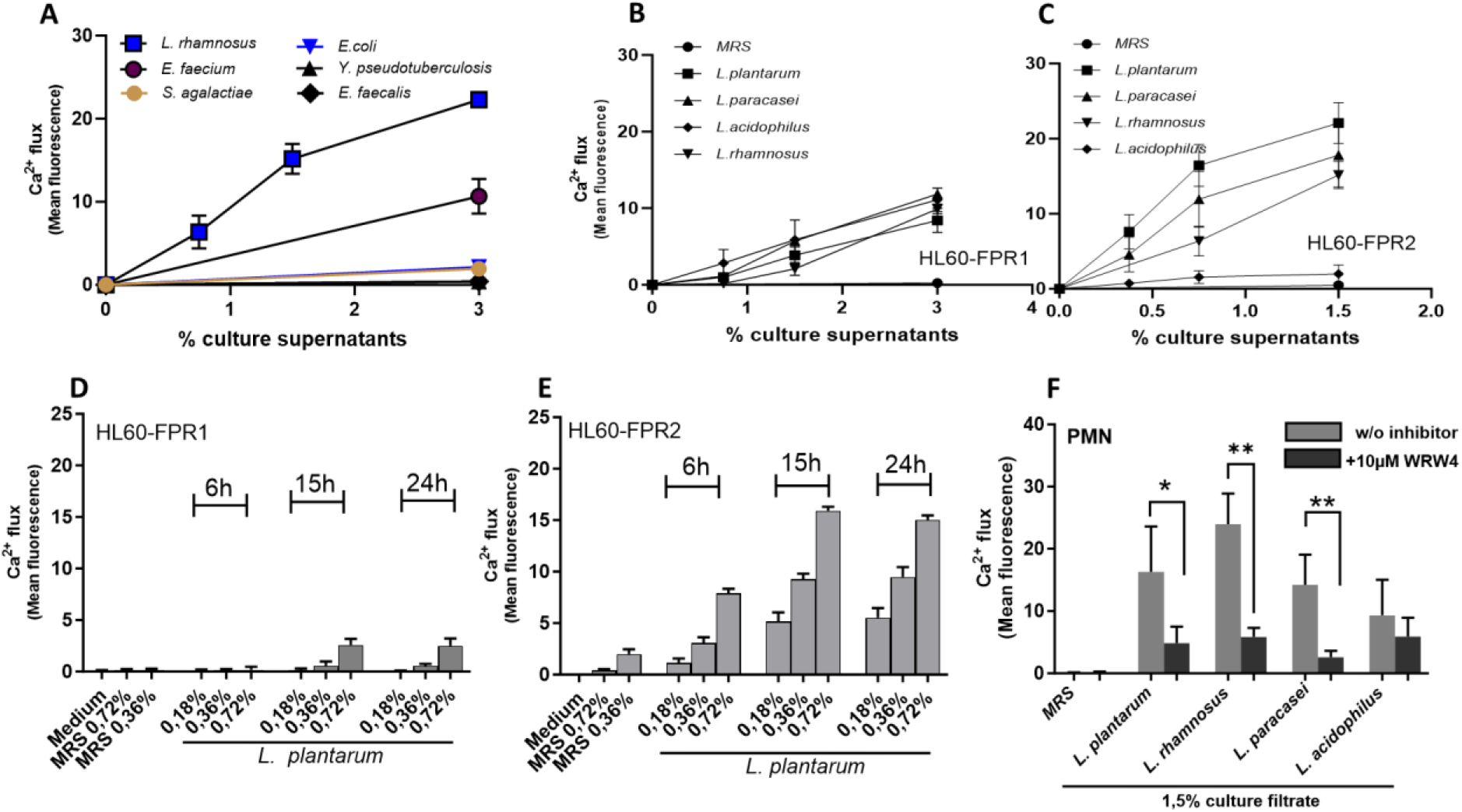
*L. plantarum, L. rhamnosus and L. paracasei* activate the formyl-peptide receptor 2. Calcium influx in FPR2-transfected HL60 cells induced by culture filtrates of different bacteria (A). Calcium influx induced by culture filtrates of the indicated lactobacilli in FPR1- (B) or FPR2- (C) overexpressing HL60 cells. Calcium influx in FPR1- (D) or FPR2- (E) overexpressing HL60 cells induced by *L. plantarum* culture filtrates cultivated for 6h, 15h or 24 h. Calcium influx in neutrophils+/- 10μM WRW4 induced by culture filtrates of the indicated lactobacilli (F). Data represent means +/-SEM of three independent experiments, and three different culture filtrates (A-C) or 2 independent experiments using two different culture filtrates (D-F) using blood from different donors (F). *P* < 0.05; \**\*, P <* 0.01, significant difference versus untreated neutrophils as calculated by paired (F). two-tailed Student’s *t* test.

### Culture filtrates of lactobacilli induce neutrophil migration, activation and enhance bacterial killing

It is known that FPR activation induces chemotaxis of neutrophils and IL-8 release. Therefore, we tested diluted culture filtrates of *L. plantarum, L. rhamnosus* and *L. paracasei* regarding their capacity to trigger migration of neutrophils and observed that all analyzed culture filtrates induced dose-dependent chemotaxis (Fig 2 A-C). Similarly, the culture filtrates of *L. plantarum, L. rhamnosus* and *L. paracasei* induced IL-8 release by neutrophils (Fig.2D). With respect to the ability of promoting neutrophil killing of bacterial pathogens, culture filtrates of all lactobacilli enhanced neutrophil killing of an important pathogen, the methicillin-resistant *Staphylococcus aureus* (MRSA) USA300 (Fig.2E-G).

**Figure 2.**
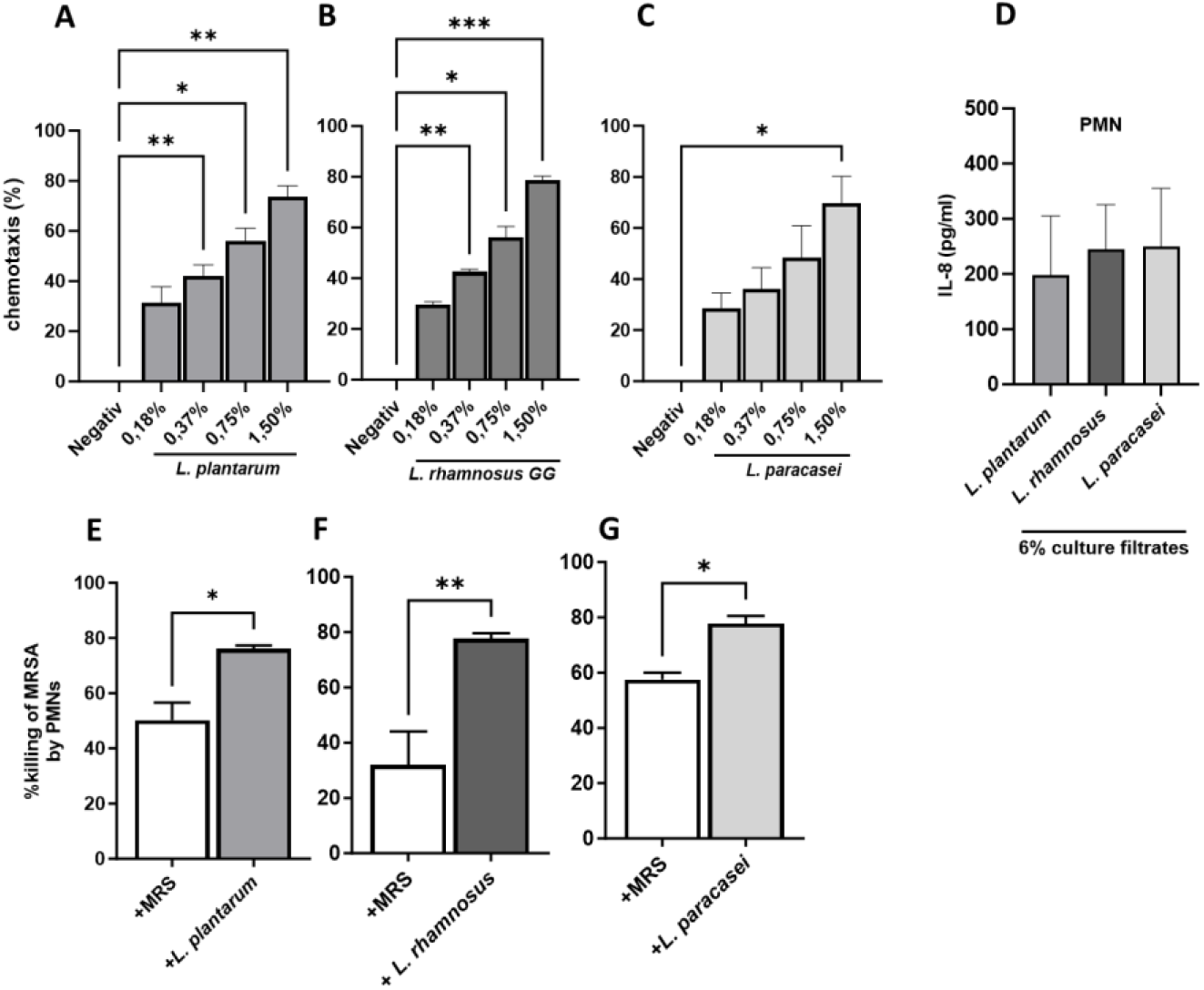
Culture filtrates of probiotic lactobacilli induce neutrophil migration, activation and enhance bacterial killing. Neutrophil migration induced by the culture filtrate of (A) *L. plantarum* (B) by *L. rhamnosus*, (C) *by L. paracasei* after 80 minutes. IL-8 release by neutrophils stimulated for 5 hours with the indicated culture filtrates of lactobacilli (E). Killing of S. *aureus* USA300 (MRSA) by neutrophils preincubated or not with culture filtrates of the indicated lactobacilli (E-G). Data represent mean+/- SEM of at least three independent experiments, using neutrophils from at least three different donors, (A-G). *P* < 0.05; \**\*, P <* 0.01 \*\**\*, P <* 0.001, significant difference versus the indicated control treated neutrophils as calculated by one-way Anova (A-C) or unpaired (E-G) two-tailed Student’s *t* test.

### The FPR2 ligands released by lactobacilli are peptides

To elucidate if the measured FPR2 activity is due to peptides present in the culture filtrates we treated culture filtrates of lactobacilli for two hours with proteinase K and analyzed afterward their capacity to activate FPR1 or FPR2. In case of the culture filtrate of *L. paracasei, L. acidophilus* and *L. rhamnosus* FPR1 activity was not influenced; only in *L. plantarum* a slight reduction of FPR1 activity was observed (Fig.3A). In contrast, proteinase K treatment markedly reduced FPR2 activity in all culture filtrates, except in the control (Fig.3B). Since FPR2 can be activated by formylated and nonformylated ligands, we tested whether inhibition of the bacterial deformylase by actinontin during growth of *L. plantarum* enhanced only FPR1 or also FPR2 activation. Inhibition of the deformylase led to enhanced FPR1 and FPR2 activity (Fig.3 C, D).

**Figure 3.**
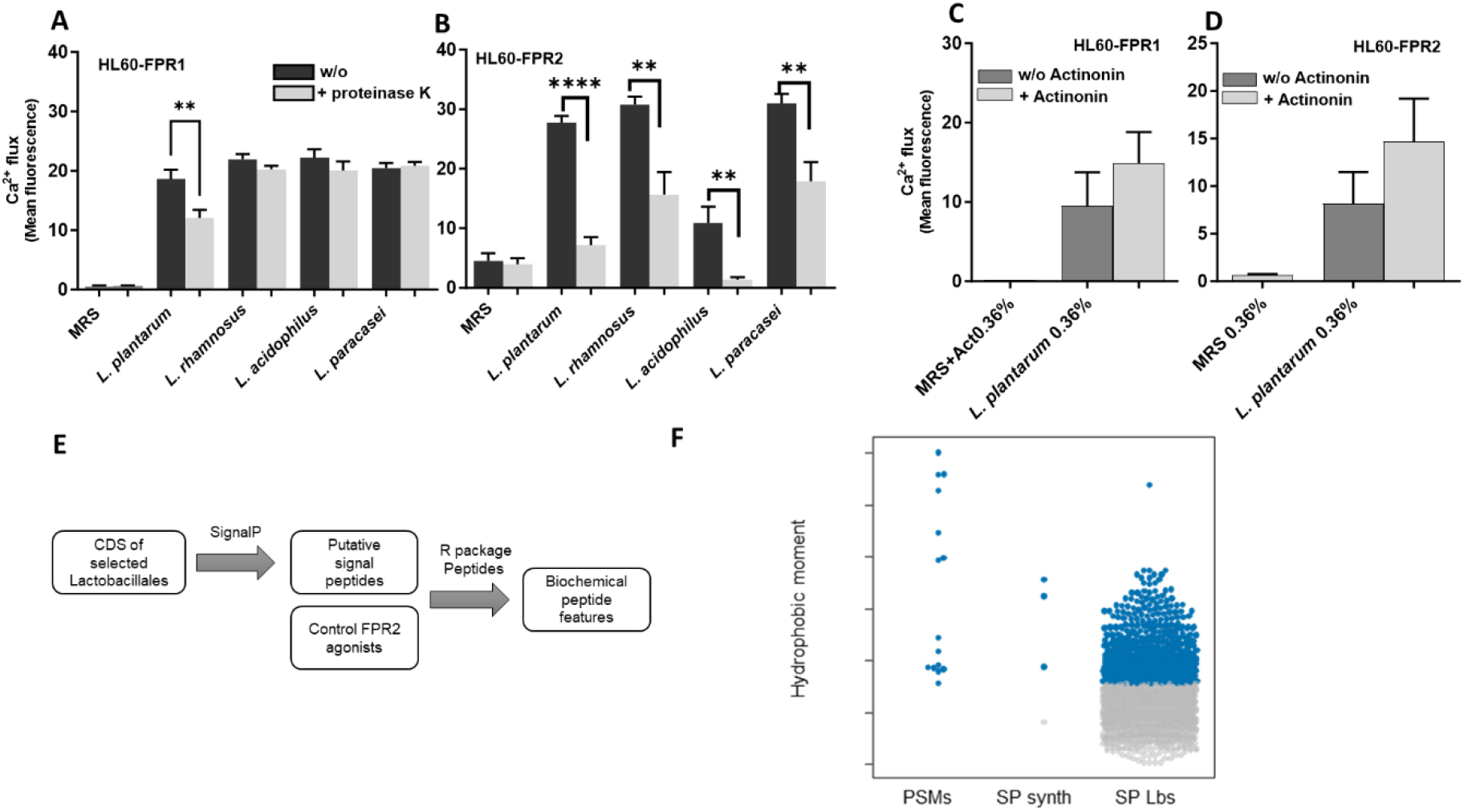
Proteinase K digestion of lactococcal culture filtrates reduces FPR2 activation. Activation of FPR1-(A) or FPR2-(B) overexpressing HL 60 with the culture filtrates (6%) digested for 2 hours +/- proteinase K (5mg/ml). Stimulation of FPR1-(C) or FPR2-(D) overexpressing HL 60 by the culture filtrates of *L. plantarum* treated or not with actinonin. Workflow for determining peptides features in silico. Potential signal peptides of publicly available or predicted coding sequences (CDS) of selected Lactobaciliales were obtained using the program SignalP. The obtained peptides and known FPR2 agonists of PSM types alpha and beta were analysed using the R package “Peptides” to obtain biochemical features (E). The hydrophobic moment as measure of FPR2 activation potential of signal peptides from lactobacilli (n=1565) is compared with PSMs (alpha and beta class is indicated) and synthetic peptides used in this study. The lowest hydrophobic moment of known FPRs2 against (PSMs) was used to determine a threshold to identify signal peptides of Lactobacillaceae with FPR2 activation potential (highlighted in blue). Data represents mean+/- SEM of three independent experiments (A, B) or two independent experiments (C, D). *P* < 0.05; \**\*, P <* 0.01 \*\**\*, P <* 0.001, \*\*\**\*, P <* 0.0001 significant difference versus corresponding non-digested culture filtrates (A, B) as calculated by unpaired two-tailed Student’s *t* test.

Amphipathic, positively charged signal peptides – and subfragments thereof – originating from the cleavage of the N-termini of bacterial proteins fulfill all the criteria for strong FPR2 agonists (Kretschmer et al. 2015). This suggests that by releasing many of such peptides, bacteria may also be potent activators of human cells via FPR2. We sequenced the genomes of the isolated lactic acid bacteria (https://www.ncbi.nlm.nih.gov/bioproject/1107177) and performed computational analysis of potential signal peptides with such properties (N-terminal formylation, a length of 18-45 amino acids, amphipathic) using predicted coding sequences from the newly sequenced genomes as well as publicly available coding sequences of different Lactobacillaceae species (table 1, Fig 3E, F). We then determined the hydrophobic moment for allobtained signal peptide as compared to a control group consisting of known FPR2 activators (of alpha and beta PSMs). We found that 624 potential subfragments (of 1565 tested Lactobacillaceae signal peptides) possessed the biochemical characteristics of potential FPR2 activators (Fig 3F).

**Table 1:** Overview of the bacterial species used for the characterization of potential signal peptides. Complete data of used peptides can be obtained from Supplementary table 1.

| Genus | Species | Strain | Source |
| --- | --- | --- | --- |
| Lactiplantibacillus | plantarum | ASM326940 | NA |
| Lactobacillus | acidophilus | Do3 | commercially available probiotics |
| Lactobacillus | acidophilus | La14 | GenBank: CP005926.2 |
| Lactocaseibacillus | paracasei | Do5 | Isolated from yoghurt |
| Lactiplantibacillus | plantarum | Do4 | commercially available probiotics |
| Lactocaseibacillus | rhamnosus | Do7 | commercially available probiotics |
| Lactobacillus | rhamnosus | GG | GenBank: GCA_003353455.1 |
| Lactococcus | acidophilus | Do6 | NA |
| Lactococcus | lactis | IO1 | GenBank: AP012281.1 |
| Lactococcus | piscium | MKFS47 | GenBank GCA_000981525.1 |
| Limosilactobacillus | reuteri | ASM918472 | NA |

### Signal peptides of *L. plantarum* strongly activate FPR2

To determine whether signal peptides of *L. plantarum* could represent potential FPR2 ligands, we synthesized some of them and analyzed their FPR2 activity. Signal peptides of *L. plantarum* predominantly induced calcium influx in FPR2-overexpressing HL60 cells. Only at high concentration of the two signal peptides ALA and KGM a slight calcium release could be observed in FPR1-overexpressing HL60 cells. Since culture filtrates of lactobacilli enhanced killing of MRSA by neutrophils, we investigated whether these peptides induced oxidative burst and promoted the expression of receptors involved in phagocytosis. All analyzed signal peptides induced oxidative burst, in particular the most active FPR2 ligands (Fig.4C). In addition, these peptides promoted dose-dependent expression of the complement receptors CD11b and CD35, and the peptides KGM as well as ALA also induced the expression of the FC receptor CD64.

**Figure 4.**
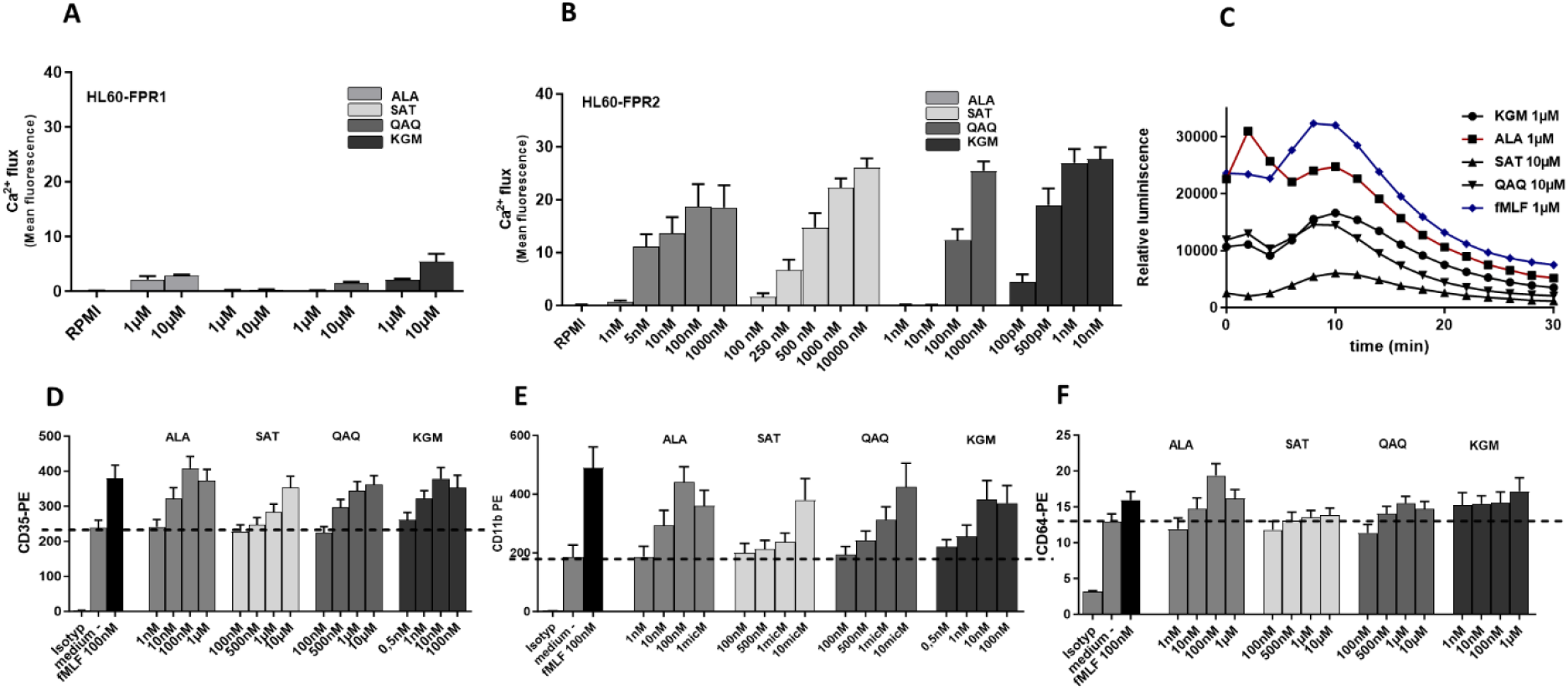
Signal peptides of *L. plantarum* strongly activates FPR2. Calcium influx induced by signal peptides (formyl-MAKFRRLVLLSLSLGLALAGGCRSPDALA (ALA), formyl-MMRGMGNMQSMMKQMKKMQAQ (QAQ), formyl-MMKITKPFRMSLVKAKGM (KGM), and formyl-MLSKSSAPKTKQQATSTKVTSKQTTNQSAT (SAT) of *L. plantarum* in FPR1 (A) or FPR2 (B)-overexpressing HL60 cells. Oxidative burst induced in neutrophils by the indicated peptides (C). Expression of complement receptor CD-35 (D), CD11b (E) or FC-receptor CD64 by neutrophils after stimulation with the indicated signal peptides. Data represent means (C) +/-SEM of three independent experiments (A, B, D-F), using neutrophils from three different blood Donors (C-E).

**Figure 5.**
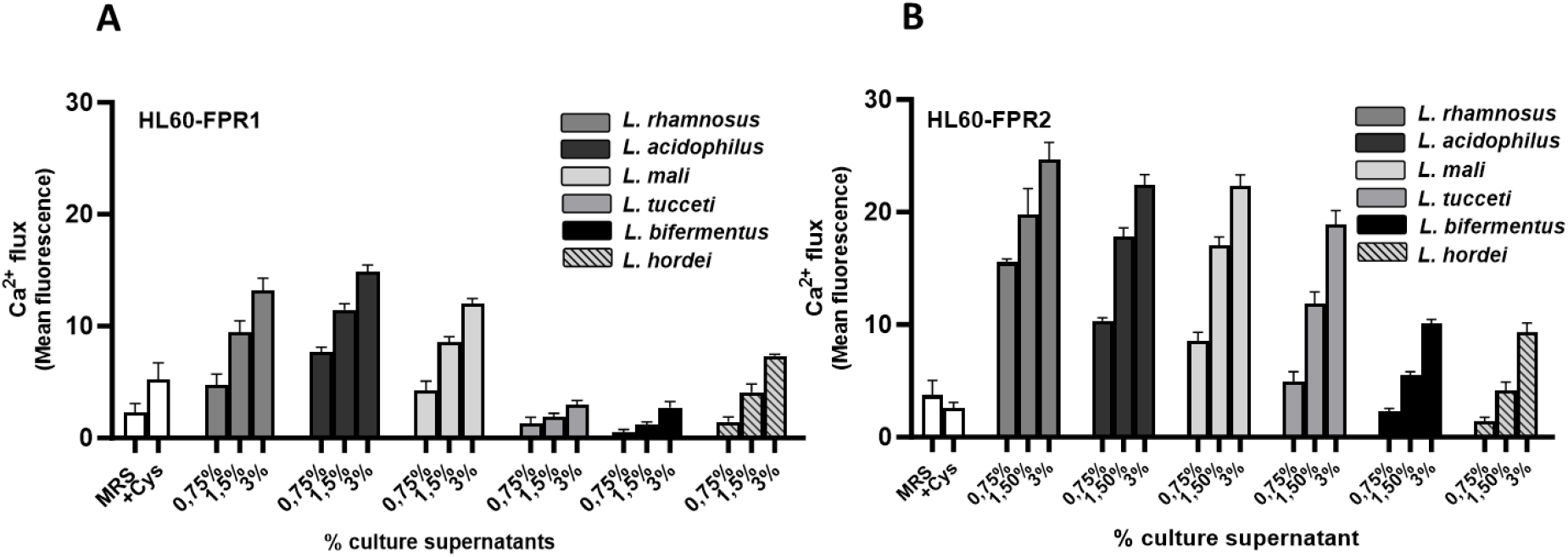
The ability to activate the formyl-peptide receptor 2 is common to many lactobacilli. Calcium influx induced by control (MRS, or MRS+0,05% cysteine) or culture filtrates of the indicated lactobacilli (*Loigolactobacillus bifermentans, Lactobacillus tucceti, Lactobacillus tucceti, Lactobacilllus acidophilus, Lactobacillus rhamnosus LGG, Liquorilactobacillus mali, Liquorilactobacillus hordei)* in FPR1 (A) or FPR2 (B)-overexpressing HL60 cells. Data represent mean and SEM of three independent experiments.

### Culture filtrates of various lactobacilli activate predominantly FPR2

To find out whether FPR2 activity is only detectable in response to some lactobacilli or whether it is a characteristic of many lactobacilli strains, we examined the ability of various culture supernatants of different lactobacilli (*L. rhamnosus, Liquorilactobacillus mali, L. acidophilus, Lactobacillus tucceti, Loigolactobacillus bifermentans*, Liquorilactobacillus hordei, Table 2) to activate FPR. Interestingly, we detected FPR2 activity to various degrees in response to all analyzed culture filtrates, whereas the FPR1 activity was rather low in some of these strains (*L. tucceti, L. bifermentus, L. hordei)*.

**Table 2:** Overview of the bacterial species used for Analyzes of FPR1 and FPR2 activation (Fig. 5).

| Genus | Species | Strain |
| --- | --- | --- |
| *Loigolactobacillus* | *bifermentans* | DSM 20003(Sun et al. 2015) |
| *Lactobacillus* | *tucceti* | DSM 20183(Sun et al. 2015) |
| *Lactobacillus* | *acidophilus* | DSM 20079(Sun et al. 2015) |
| *Lactobacillus* | *rhamnosus* | LGG |
| *Liquorilactobacillus* | *mali* | ATCC 27304 |
| *Liquorilactobacillus* | *hordei* | DSM19519 (Rouse, Canchaya, and van Sinderen 2008) |

## Discussion

Bacterial metabolites and MAMPs are sensed by a variety of mammalian receptors, thereby initiating inflammatory responses, eliciting leukocyte chemotaxis, or altering cell proliferation and homeostasis. The best known bacterial FPR2 ligands are the phenol-soluble modulins (PSMs), formylated peptide toxins which are encoded in the core genome of pathogenic and commensal staphylococci (Rautenberg et al. 2011). In addition, host-derived peptides like the cathelicidin LL37 or lipids like resolvin D1 and lipoxin A4 have been shown to be associated with inflammation and bind to FPR2 (Weiss and Kretschmer 2018). These indicate that FPR2 can recognize a diverse set of ligands. In the current work we demonstrated that various lactobacilli release large amounts of ligands that can specifically activate FPR2 and possibly mediate their probiotic effects. We found that among the strongest FPR2 activators are lactobacilli that were reported to promote wound healing and prevent inflammation in the gut (Kelm and Anger 2022).

Bufe et al. showed that signal peptides provide a large pool of FPR agonists with different amino acid sequences but a conserved secondary structure (Bufe et al. 2015). Mass spectrometry analyses of bacterial secretomes found complete signal peptides as well as N-terminal fragments in culture filtrates, which show that these molecules can be secreted by bacteria (de Souza et al. 2011; Ravipaty and Reilly 2010). It has been speculated that the release occurs via lysis or autolysis of the bacteria.

However, also the release via membrane vesicles could be an explanation which is supported by vesicle releasing *L. plantarum* (Yu et al. 2022). Interestingly, systematic analysis of the signal peptides and culture filtrates of different lactobacilli suggested that further bacterial species from the genera Lactobacillus, Akkermansia, and Enterococcus with described probiotic potential (Li et al. 2024; Im et al. 2023) release putative FPR2 ligands.

Using synthetic signal peptides of *L. plantarum*, we could show that they induce ROS in neutrophils. It is known that FPR ligands can induce ROS in a receptor-dependent manner not only in neutrophils via NADPH oxidase (NOX)2 but also in gut epithelia cells via NOX1 (Leoni et al. 2013). It has been described that *L. rhamnosus* GG stimulated myenteric production of ROS in mice via FPR1. However, the authors of the same study also observed reduced ROS production in myenteric ganglions of FPR2 knockout cells indicating that FPR2 is involved in ROS production as well (Chandrasekharan et al. 2019). Such ROS production plays a role in epithelia cell proliferation, migration, barrier function and wound healing in the gut (Alam et al. 2016; Jones et al. 2013). Especially in case of intestinal wound healing, FPR2 seem to play a crucial role because decreased numbers of infiltrating monocytes were observed in healing wounds of FPR2/3 knock out mice (Birkl et al. 2019).

A rare sequence variant in NOX1 is responsible for pediatric onset of irritable bowel disease (IBD) which highlights the involvement of human NOX1 in regulating wound healing by altering epithelial cytoskeletal dynamics at the leading edge and directing cell migration (Khoshnevisan et al. 2020). Furthermore, colonic epithelial cells in FPR2-deficient mice displayed defects in commensal bacterium-dependent homeostasis leading to shortened colonic crypts, reduced acute inflammatory responses, delayed mucosal restoration after injury, and increased azoxymethane-induced tumorigenesis (Chen et al. 2013). Moreover, it has been shown that mucosal expression of FPR2/ALX mRNA is 7-fold increased in the gut of patients with ulcerative colitis (Vong et al. 2012). Fpr2 deficiency increased susceptibility to chemically induced colitis, delayed the repair of damaged colon epithelial cells, and heightened inflammatory responses. Additionally, the population of *Escherichia coli* was observed to increase in the colon of Fpr2 ^−/−^ mice with colitis (Chen et al. 2023). These results indicate that FPR2 is critical in mediating homeostasis, inflammation, and epithelial repair processes in the colon. It is tempting to speculate that FPR2 ligand-producing lactobacilli may positively influence the outcome of inflammatory gut diseases in a FPR2 dependent manner.

## Material and Methods

### Bacteria and Cell Lines

*Lacticaseibacillus rhamnosus, Lactiplantibacillus plantarum, Lacticaseibacillus paracasei* were isolated from probiotic compounds. A list of the strains used is provided in Table 1. Bacterial culture supernatants were obtained from cultures grown in MRS for 48h at 37° under low oxygen conditions in closed Falcon tubes without agitation. Bacterial cultures were centrifugated and supernatants subsequent filtrated through 0.2-μm pore size filters (Merck). HL60 cells stably expressing human FPR1, FPR2, have been described recently (Christophe et al., 2001; Dahlgren et al., 2000). These cells were grown in RPMI medium (Biochrom) supplemented with 10% FCS (Sigma-Aldrich), 20 mM Hepes (Biochrom), penicillin (100 units/ml), streptomycin (100 μg/ml) (Gibco), and 1 x Glutamax (Gibco). Transfected cells were grown in the presence of G418 (Biochrom) at a final concentration of 1 mg/ml. Culture filtrates from further Lactobaciliales(Table 2); L. *bifermentus, L. tucceri, L. rhamnosus, L. mali, L. horderi* were prepared in MRS or MRS+0,05% Cysteine for *L. acidophilus. S. aureus* USA300 lac (Wang et al. 2007) was cultivated overnight in Tryptic Soy Broth.

### Peptides

Signal peptides from *Lactiplantibacillus plantarum*, namely, formyl-MAKFRRLVLLSLSLGLALAGGCRSPDALA (ALA), formyl-MMRGMGNMQSMMKQMKKMQAQ (QAQ), formyl-MMKITKPFRMSLVKAKGM (KGM), and formyl-MLSKSSAPKTKQQATSTKVTSKQTTNQSAT (SAT), were synthesized by EMC Tübingen. fMLF was purchased from Sigma Aldrich.

### Sequencing of Lactobacilli

The quality of the raw reads was assessed using FastQC (v0.11.8). Trimming of the raw reads, whole genome assembly, and refining the assembly were carried out using Trimmomatic (v0.39), SPAdes (v 3.15.5), and Pilon (v1.24), respectively (Bolger, Lohse, and Usadel 2014; Prjibelski et al. 2020; Walker et al. 2014). These steps were performed within the Shovill pipeline. Finally, Prokka (v1.14.6) was utilized for the functional annotation of the assembled genomes (Seemann 2014).

### Isolation of human neutrophils and Chemotaxis

Human neutrophils were isolated by standard Ficoll/Histopaque gradient centrifugation (Dürr et al., 2006). Blood was kindly donated by healthy volunteers (age 20–50) upon informed consent. The studies were approved by local medical ethical committee (reference numbers 750/2018BO2, 054/2017BO2). For the analysis of the chemotactic capacities of neutrophils exposed to culture filtrates, neutrophils were loaded with 3 μM 2′,7′-bis-(2-carboxyethyl)-5-(and-6)-carboxyfluorescein, acetoxymethyl ester (BCECF-AM, Molecular Probes). The migration along gradients of the indicated stimuli was monitored using 3-μm polycarbone trans-well membranes (Greiner) (Schlatterer et al. 2021). The compartments below the cell culture inserts contained the diluted supernatants. After 80 minutes, the cell culture inserts were removed, and the fluorescence intensity of the migrated neutrophils in the lower compartments was measured using a BMG Labtech CLARIOstar plate reader.

### Measurement of Calcium Ion Fluxes in Human Neutrophils and HL60 Cells

Calcium fluxes were analyzed by stimulating cells loaded with Fluo-3-AM (Molecular Probes) and monitoring fluorescence with a FACS Calibur flow cytometer (Becton Dickinson) as described recently (). For measuring the influence of WRW4 1 x 10^6^ cells/ml were preincubated with WRW4 at a final concentration of 10μM, for 20 min at room temperature under agitation. To stimulate neutrophils and HL60 cells, peptides were used at the indicated concentrations in RPMI + HSA0,05%. Culture supernatants were used at indicated dilutions in RPMI +HSA 0,05%. Measurements of 2,000 events were performed and calcium flux was expressed as relative fluorescence. To elucidate if FPR2 -activating compounds in Lactobacilli culture filtrates are of proteinaceous nature, culture filtrates (pH neutralized) were treated with proteinase K beads (1mg/ml) and incubated for 1 h at 37°C under agitation. Proteinase K Eupergit H C beads were subsequently removed by centrifugation (10 min at 250xg). For inhibition of deformylase with actinonin culture filtrates were obtained from *L. plantarum* grown 24 hours in MRS at 30°C in the presence or absence of 10μg/ml actinonin. Bacteria were removed by centrifugation and supernatants were passed through a 0.22 mm-pore size sterile filter. Proteolytically digested culture filtrates or Actinonin treated culture filtrates were used in the calcium flux assay with FPR2 -transfected HL60 cells as described above.

### IL-8 detection

The release of IL-8 from neutrophils was measured with a human IL-8/CXCL8 ELISA Kit (R&D). Primary neutrophils were stimulated with the indicated dilutions of sterile filtered culture filtrates of lactobacilli for 5 hours. Human IL-8 detection in the cellular supernatant was performed according to the IL-8 ELISA vendor’s manual using FluoStar optima.

### Killing Assay

*S. aureus* USA300 was inoculated at OD 600 0.1 in tryptic soy broth (TSB) or and grown for 4 h under aerobic conditions (medium to flask ratio 1:5) followed by three washing steps with PBS. For optimal recognition by neutrophils, bacteria were opsonized with 10% pooled normal human serum (NHS) for 60 min and neutrophils and bacteria and were used at a MOI of 0,1. For the bacterial killing assay in the presence of culture filtrates, the neutrophils and bacteria were seeded in a 24-well plate at an MOI of 0.1 and incubated for 60 minutes with 50μl culture filtrates or MRS diluted in PMN media (1:10). After 1 hour, 100 μL of each sample was collected, and the neutrophils were lysed with ddH2O for 15 minutes at 4°C,1000 rpm. Serial dilutions of the samples were plated on TSA plates using an IUL EDDY Jet 2 spiral plater. On the following day, the colony-forming units (CFUs) were counted with anIUL Flash & Go instrument.

### Expression of Complement Receptor and FCγ Receptor

Neutrophils were seeded into a 96-well-plate and stimulated with ALA, KGM, QAQ and SAT peptides at the indicated concentrations for 1 hour. Subsequently, the supernatant was discarded, and then neutrophils were incubated with PE-labeled antibodies against CD11b (BD Pharmingen), CD35 (Miltenyi Biotech), CD64 (Miltenyi Biotech), or an IgG isotype control (Miltenyi Biotech) for 30 minutes on ice. Next, the neutrophils were fixed with 3.7% formaldehyde, and the fluorescence intensity of the neutrophils was determined using a BD FACSCalibur instrument, and the mean fluorescence intensity was analyzed.

### Analysis of signal peptides

Signal peptides of bacterial organisms were identified in publicly available protein sequences (source: NCBI datasets) from selected strains (see Table 1) using the bioinformatic tool SignalP (Version 5.0)(Almagro Armenteros et al. 2019). As positive control for FPR2 agonists we utilized well-defined alpha- and beta-PSMs from staphylococcal species with amphoteric characteristics. Biochemical peptide features were determined using the R package Peptides {Osorio, 2015 #1510} (https://doi.org/10.32614/RJ-2015-001). To calculate the amphipathic peptide features according to the Eisenberg method {Eisenberg, 1984 #1509} we utilized the provided “hmoment” function. Peptide features were analyzed, and figures were generated using R studio (version 4.2.3).

### Oxidative burst measurements

Reactive oxygen release (ROS) by human neutrophils was measured over a time of 30 minutes by monitoring luminol-amplified chemiluminescence using 282 μM luminol (Sigma Aldrich). Neutrophils were stimulated either with negative control (Hank Balanced Salt Solution, Sigma Aldrich) or with fMLF as positive control (1μM), KGM (1μM), QAQ (10μM), SAT (10μM), or ALA (1μM). Luminescence was measured using Fluostar optima (BMG Labtech).

### Statistics

All statistical analyses were performed with Graph Pad Prism 10.0 (GraphPad Software, La Jolla, USA). An unpaired or paired two-tailed Student’s t test was performed to compare two data groups, while more than two data groups were analysed by one-way ANOVA with Dunnett’s multiple comparisons test, if not otherwise noted.

## Conflict of Interest

The authors declare that the research was conducted in the absence of any commercial or financial relationships that could be construed as a potential conflict of interest.

## Author Contributions

D.K., R.R. and D.G. designed the experiments; D.K., A.E., R.R., D.G. and P. W. O’Toole performed the experiments; D.G, B.K, A.P., and D.K. edited the manuscript and interpreted the data.

## Funding

This study was funded by grants from the German Research Foundation. A.P. is supported by the Cluster of Excellence EXC 2124 ‘Controlling Microbes to Fight Infections’ project ID 390838134

## Acknowledgments

We thank Cosima Hirt and Cordula Geckeler for excellent technical support, and Libera Lo Presti for critical reading/editing the manuscript. The authors acknowledge support by the High Performance and Cloud Computing Group at the Zentrum für Datenverarbeitung of the University of Tübingen, the state of Baden-Württemberg through bwHPC and the German Research Foundation (DFG) through grant no INST 37/935-1 FUGG.

## Notes

https://www.ncbi.nlm.nih.gov/bioproject/1107177

### Competing Interest Statement

The authors have declared no competing interest.

